# Comprehensive surveys of endangered western hoolock gibbons (*Hoolock hoolock*) in Bangladesh more than double the country’s total population estimate

**DOI:** 10.1101/2025.03.13.642956

**Authors:** M Tarik Kabir, Susan Lappan, Shahrul Anuar Mohd Sah, Nadine Ruppert

## Abstract

The western hoolock gibbon (*Hoolock hoolock*, Harlan 1834) is a Critically Endangered primate species in Bangladesh and is globally Endangered, yet its distribution across the highly anthropogenic and disturbed landscapes of southeast and northeast Bangladesh is not well understood. We assessed gibbon occurrence in habitats across Bangladesh through community engagement activities from January 2018 to June 2022 and conducted gibbon population surveys in the field from January 2019 to March 2024 using the total count method to estimate abundance in most habitats. Our results confirmed gibbon occurrence in 46 locations in 42 forest fragments in southeast and northeast Bangladesh and we estimate the current gibbon population in Bangladesh to include ≥816 individuals in ≥254 groups. Only 10 of the 42 fragments where gibbons occurred supported more than 20 individuals. Gibbons in most of these habitats are at extremely high risk of local extinction due to very small population sizes and ongoing habitat destruction, hunting, and agricultural expansion. Urgent conservation actions, including community-based conservation to combat hunting and illegal trade and to prevent further agricultural encroachment, support for monitoring and enforcement of existing laws and regulations, habitat protection, and habitat restoration programs focused on landscape-level connectivity within Bangladesh and across the Bangladesh-India border, are required to mitigate threats to the remaining gibbons in Bangladesh.

## Introduction

Gibbons or small apes (family Hylobatidae) are distributed in mixed-evergreen and tropical rainforests in South and Southeast Asia (Chivers, 1977; Geissmann, 1995). The western hoolock gibbon (*Hoolock hoolock*) is distributed across northeast India, northwestern Myanmar, and eastern Bangladesh (Brockelman et al., 2019), and is the only small ape species in Bangladesh and India (Tribedi et al., 2021). The species is listed as Endangered globally due to habitat loss and hunting (Brockelman et al., 2019). Northeast India may support ≥12,000 individuals (Geissmann et al., 2013), while Myanmar is likely home to the largest population of this species but most gibbon habitats there have not yet been surveyed (Brockelman et al., 2019). The conservation situation for western hoolocks in Bangladesh, which forms much of the westernmost occurrence of gibbons globally, is dire. Western hoolocks are Critically Endangered in Bangladesh, where only small fragments of hoolock habitat remain in mixed-evergreen hill forests (Feeroz et al., 2015). However, a comprehensive assessment of the status of western hoolocks in Bangladesh is not yet available, which makes it difficult to formulate comprehensive conservation plans for the species on the global or national level.

Bangladesh occupies 147,570 km^2^ of land with different types of ecosystems. Only the mixed- evergreen hill forests of Chattogram, Chattogram Hill Tracts, Cox’s Bazar, and Sylhet region (Nishat et al., 2002) are appropriate habitat for gibbons. These forests are fragmented, with some relatively large patches still existing in the Chattogram Hill Tracts of southeast Bangladesh (Khan, 2008). These forests support many globally threatened species including Asian elephants (*Elephas maximus*), Asiatic black bears (*Ursus thibetanus*), Phayre’s langurs (*Trachypithecus phayrei*), sambar deer (*Rusa unicolor*), wild dogs (*Cuon alpinus*), Indian leopards (*Panthera pardus pardus)*, clouded leopards (*Neofelis nebulosa*), and western hoolock gibbons (*Hoolock hoolock*; UCN Bangladesh, 2015). Threats to these forests include illegal logging, encroachment, excessive resource harvesting, wildlife poaching, and smuggling, agricultural expansions inside the forested areas, and the introduction of exotic and invasive species (Chowdhury et al., 2014; Reza and Hasan, 2019). Within these landscapes, the distribution of gibbon habitats is not homogenous or contiguous but restricted to several highly degraded forests (Islam et al., 2006; FD, 2020).

Despite its status as Bangladesh’s only ape and the threats that it faces, the western hoolock gibbon has not received much attention in Bangladesh, resulting in inadequate long-term monitoring of its population. Almost 20 years ago, Islam et al. (2006) conducted a population assessment at 35 sites and estimated that a population of only about 282 western hoolock gibbon individuals remained in Bangladesh. More recently, Naher et al. (2021) surveyed 22 field sites and estimated that 469 individuals remained in eastern Bangladesh. However, neither of these estimates were based on comprehensive surveys across all potential habitat fragments, and both teams were unable to confirm the presence of gibbons at many suspected gibbon habitat patches in the northeast and southeast mixed evergreen hill forests. Rapid surveys by teams of scientists without detailed local information may not be adequate to confirm gibbon absence, as gibbons are often difficult to detect visually, especially when the terrain is challenging (Geissmann et al., 2013). A more comprehensive assessment of wildlife distribution in large landscapes can be aided by involving local communities that live in shared habitats (Shirk et al., 2012; Sun et al., 2021). In the current study, we aimed to identify new gibbon habitats in Bangladesh, to document occurrence across all known or suspected habitats, and to estimate gibbon population size and conservation status by pairing local knowledge with extensive and long-term field investigation.

## Methods

### Study areas

Bangladesh forms part of the transition zone for flora and fauna between the Indo-Himalayan and Indo-Chinese subregions (FD, 2020). Forested areas in Bangladesh consist of mixed- evergreen forests (hill forest), deciduous forests (sal forest), mangrove forests, swamp forests, and village forests (https://bforest.gov.bd/). The total forested area in Bangladesh is 1.429 million hectares, which constitutes 11% of the total land area of the country (FAO, 2015).

Gibbons are believed to occur only in the mixed evergreen hill forests of Chattogram, Cox’s Bazar and Chattogram Hill Tracts (Rangamati, Bandarbans and Khagrachari district) of southeast Bangladesh and Habiganj and Moulvibazar district of northeast Bangladesh (Figs.1 and 2).

Hill forests, which are managed by the Forest Department, comprise about 676,000 ha or 44% of the total forested area in Bangladesh (FD, 2020). Hill forests support natural terrestrial vegetation with evergreen and deciduous trees in association with bamboo groves. Major tree species include *Antidesma ghasembilla*, *Ardisia solanacea*, *Bombax* sp., *Dipterocarpus* sp. *Ficus* sp. *Tectona grandis*, *Lagerstroemia parviflora*, *Syzygium* sp., *Swintonia floribunda* (Islam et al., 2006). Tree cover in these forests ranges from 10-100% and tree height ranges from 5 m to 35 m.

Data collection

### Identification of new gibbon habitats

New gibbon habitats were identified through extensive community engagement activities in the form of consultation programs, focus group discussions (FGD), and in-person interviews with local communities, frontline staff of the local Forest Department, and wildlife enthusiasts as well as by surveying social media and newspaper posts and field surveys, and gibbon presence in these habitats was confirmed during field work from January 2018 to December 2022 (Bernard, 2006). More than 950 respondents were involved in community engagement activities (Range = 1 to 96 participants per site, mean = 27.51, SD= ±25.76). Community consultation programs were part of a broader set of community engagement activities with involvement of the Forest Department and local community members for raising awareness about western hoolock gibbons and their conservation needs. Participants for FGD were selected from the nearest villages or human settlements around forested areas in northeast and southeast Bangladesh. Wildlife enthusiasts that we interviewed comprised wildlife biologists, nature lovers, bird watchers, and ecotourists who had visited hill forests in Bangladesh. Social media and newspaper posts were closely monitored for mentions of gibbons, including observations, photographs, awareness-raising activities, and reports of rescues from the illegal trade.

### Population census

We conducted a comprehensive population census of the western hoolock gibbon using the total count method (Kabir et al., 2021) in northeast and southeast Bangladesh from January 2018 to March 2024, surveying a total area of ca. 1975 km^2^. A total of 952 days were spent in the field by the first author, TK, (411 days) and trained research assistants (541 days). In highly degraded habitats, like most in Bangladesh, low gibbon group densities and hunting can impact gibbon singing behavior; thus, supplementing auditory identification with direct visual counts is the best method to census gibbon populations in these human-dominated habitats (Yin et al., 2016; Kabir et al., 2021).

Survey sites were chosen through the results of the community engagement activities, direct field observations, social media and newspapers, and secondary information such as published and unpublished articles (i.e., scientific articles, grey literature, and dissertations). The distribution of gibbon populations in Bangladesh is patchy, and populations are mostly restricted to patches of forest in eastern Bangladesh. Each forest beat (a local management unit under the Forest Department), was considered a distinct location for the purposes of our census except.

Lawachara National Park (NP), Sangu-Matamuhuri Reserved Forest (RF), Kassalang RF because management practices and threats varied depending on the beat (Islam et al., 2006). Privately owned forests and unclassified state forests were also considered as distinct locations. Extensive fieldwork was not possible at Kassalong RF, Sangu-Matamuhuri RF, and a few other sites in southeast Bangladesh due to security problems related to the presence of insurgents. Sangu- Matamuhuri RF (Ca. 549 km^2^) and Kassalong RF (ca. 502 km^2^) are the largest hill forested areas in Bangladesh but little information is available about their wildlife (Creative Conservation Alliance, 2016). Limited fieldwork activities were carried out and trained local research assistants were engaged to collect population information for gibbons at these habitats to the extent possible.

At the beginning of the population census, gibbon groups were detected at dedicated listening posts (Brockelman and Ali, 1987; Brockelman et al., 2009). At least one observer at one listening post carefully noted the singing times and durations, compass bearings, and the total number of heard groups. Later in the day or on subsequent days, each group was located and GPS locations of the gibbon groups, or their approximate singing location, were recorded.

Whenever possible, detected groups were visually observed and the group compositions were recorded. Gibbons were categorized as adult males, adult females, subadults, juveniles, or infants by their coat colors, body size, and behavior (Ahsan, 1994; Kakati, 2009). Neighboring gibbon groups were distinguished during the auditory survey if visually contacted or by location (songs produced at least 500 m apart were considered to have been produced by different groups), group composition, and singing times (songs produced in different locations at the same time were considered to have been produced by different groups). Then, all gibbon groups identified in each habitat (except in some locations in Chattogram Hill Tracts) were given a unique identification number, and each of these groups was monitored for at least 10 days to obtain precise information on the group composition and to map the territorial boundaries between neighboring groups to avoid double counting. The gibbon populations of Patharia Hill RF (Lathitila), Rajkhandi RF (Korma), Atora Hill RF (Ragna and Sagornal) in northeast Bangladesh and Dhopachari Forest Belt (Dhopachari), Baishari-Bangdhepa WS and Sheikh Jamal Inani NP in southeast Bangladesh were each monitored for a period of at least a year, with observations on for 3 to 5 consecutive days each month with support from trained local research assistants.

A total ca. 1975 km^2^ were surveyed across the forested areas where gibbon occurrence was suspected based on the initial habitat identification phase. Gibbon habitats were considered as occupied if the presence of the gibbon were confirmed during the field surveysThe total gibbon population was estimated by compiling the total group count and the assessed groups sizes for groups from which group sizes were available. Mean group sizes for each population were used to estimate the number of individuals in groups contacted in the surveys for which group size could not be accurately assessed for all groups within a habitat. Group size assessment was not possible in certain areas in southeast Bangladesh (Kaptaimukh, Panchari, Lemuchari, Kurokkhong, Alikhong, Bolipara); the total number of groups in these habitats was estimated from gibbon calling locations, using the methods described above, and the number of individuals in these populations was conservatively estimated by multiplying the number of groups by the lowest mean group size from sample of populations, 3 individuals/group. Solitary individuals could be distinguished from pairs in visual surveys and in vocalization surveys due to the absence of duetting behaviour. Solitary individuals were also considered as distinct groups for the total group count, but group sizes of 1 were not included in mean group size estimates.

Gibbon population densities were calculated using both the total area of the habitat and the total forested area within the surveyed area to account for the fact that not all of the land within a fragment comprised suitable gibbon habitat.

### Threat assessment

The severity of habitat destruction through illegal felling, hunting, and ongoing expansion of agricultural activities was estimated at each habitat site to develop a matrix of threats to the gibbons through direct visual field observations and via questionnaires distributed during interviews with informants (Table 1). Habitat destruction due to illegal tree felling and habitat destruction due to agricultural activities were each scored from 1 to 5 (minimal to severe). As hunting immediately affects gibbon populations, which can lead to rapid local extinctions, it was scored on a scale from 1 to 10 (low to very high). Finally, habitats were categorized using the sum of these three scores, resulting in a scale of 3 to 20, as “Highly Threatened” (15 to 20), “Medium Threatened” (9 to 14), and “Slightly Threatened” (5 to 8) (Table 1).

**Table 1:** Threats assessment matrix of the gibbon habitats of Bangladesh.

| **Habitat Destruction by direct visual observations** |  |  |
| --- | --- | --- |
| **Score** | **Intensity** | **Criteria** |
| 5-4 | High | Sign of vehicles for mass transportation of firewood and timber wood collected inside the forests or direct observation of the mass illegal resources harvesting during field work. |
| 4-3 | Medium | Commercial gardening inside the forests and frequent harvesting of timber for commercial use. |
| 2-1 | Low | Timber wood collection for self and limited commercial purposes; Almost no commercial gardening inside the forest |
| **Hunting** |  |  |
| 10-7 | Highly Threatened | Confirmation of gibbon hunting or illegal gibbon trade through community engagement activities, interviews with local communities, and information from local research assistants. |
| 6-4 | Threatened | Local communities heard or confirmed hunting or illegal trade involving primates hunting or illegal trade as well as wildlife hunting. |
| 3-1 | Slightly Threatened | Almost no wildlife hunting. |
| **Agricultural Activities including orchards** |  |  |
| 5-4 | Highly destructive | Expansion of Khasia Punji (indigenous Khasia community village), Jhoom, and betel leaf vineyard into the forested areas on a large scale. |
| 4-3 | Medium Destructive | Lemon orchards, limited-scale betel leaf vineyards plantation and other agricultural practices are frequently seen. |
| 2-1 | Slightly destructive | Lemon orchards and other agricultural activities are scattered in very small areas. |

| **Matrix** |  |  |
| --- | --- | --- |
| **Score** | **Color** | **Impacts** |
| 15 to 20 |  | Gibbons are threatened by multiple intense anthropogenic factors simultaneously and we estimate that the risk of local extinction is very high if effective conservation action is not implemented (**Highly** |
| 9 to 15 |  | Gibbons face one or more threats of moderate intensity, and the population is at risk of local extinction if threats are not addressed (**Medium Threatened**) |
| 5 to 8 |  | Some external threats to gibbons and gibbon habitat were detected. Conservation action to reduce these threats and population monitoring are recommended (**Slightly Threatened**) |

## Results

A total of 51 habitats were identified as possible gibbon habitats during the first phase of the project. More than half of these (N=35, or 68.6% of habitats) were identified through community engagement activities, and the remainder were identified through field observations (7.8%) or secondary information (23.5%).

The occurrence of the gibbons in these habitats was confirmed in 46 out of 51 habitats during the subsequent field investigations (Table 2), including 26 habitats in southeast Bangladesh and 20 in northeast Bangladesh. Community members also suggested that gibbons occur in Bhaginar Jhiri, Tonkwabati, Rajghat, and Bomarighona but despite extensive survey effort, we did not detect gibbons at those sites. Previous surveys reported the presence of gibbons at Bamu Reserved Forest (Islam et al., 2006) but we did not find gibbons there in our field observations. The 46 confirmed gibbon habitats are administered by nine forest divisions of the Bangladesh Forest Department in northeast and southeast Bangladesh (Table 2) and gibbons were found to occur in protected areas (n = 11), reserved forests (n = 22), unclassified state forests (n = 8), and privately owned forests (tea estates, n = 4; settlement Khasia Punji, n =1). The 46 occupied habitats are located in 42 forests since some forest fragments span multiples Forest Beats. Confirmed gibbon habitats ranged from 0.13 km^2^ to 549.51 km^2^ in size (n = 37, mean ± SD = 36.98 ± 115.66 km^2^), but most habitats were <10 km^2^ in size (25^th^ percentile = 1.27 km^2^, 75^th^ percentile = 9.70 km^2^). We consider the Kassalong Reserved Forest and Sangu-Matamuhuri Reserved Forest as a single forest fragment each, since we could not categorized our results from Kassalong Reserved Forest and Sangu-Matamuhuri Reserved Forest according to Forest Beat jurisdiction.

Our surveys showed that the estimated gibbon population of Bangladesh consists of at least 816 individuals in 254 groups, with at least 458 individuals in 148 groups occurring in southeast Bangladesh and 358 individuals in 106 groups in northeast Bangladesh. The presence of 25 additional groups that were detected during auditory surveys or reported by local research assistants could not be confirmed during the ground-truthing phase; these groups were not included in the total population estimate.

Table 2: Population assessment of western hoolock gibbons at southeast and northeast Bangladesh This table will be available on request.

Of the 42 forest fragments that were confirmed to be occupied by gibbons, over half of the habitats (n=23) supported ≤10 individuals, while nine fragments supported 11 to 20 individuals, and ten fragments had populations including >20 individuals.

Gibbons group sizes ranged from two to six individuals with a mean ± SD group size of 3.34±1.09 individuals (n = 108). Group sizes in southeast Bangladesh (N = 53, mean ± SD = 3.22±1.12) were similar to those in northeast Bangladesh (n = 76, mean ± SD = 3.42±1.07; range 2 to 6 (Fig. 3). Group sizes were somewhat lower in completely isolated gibbon habitats that supported <3 groups (n = 23, mean ± SD group size = 3.09 ±1.06 individuals; range = 2 to 5).

**Figure 1:**
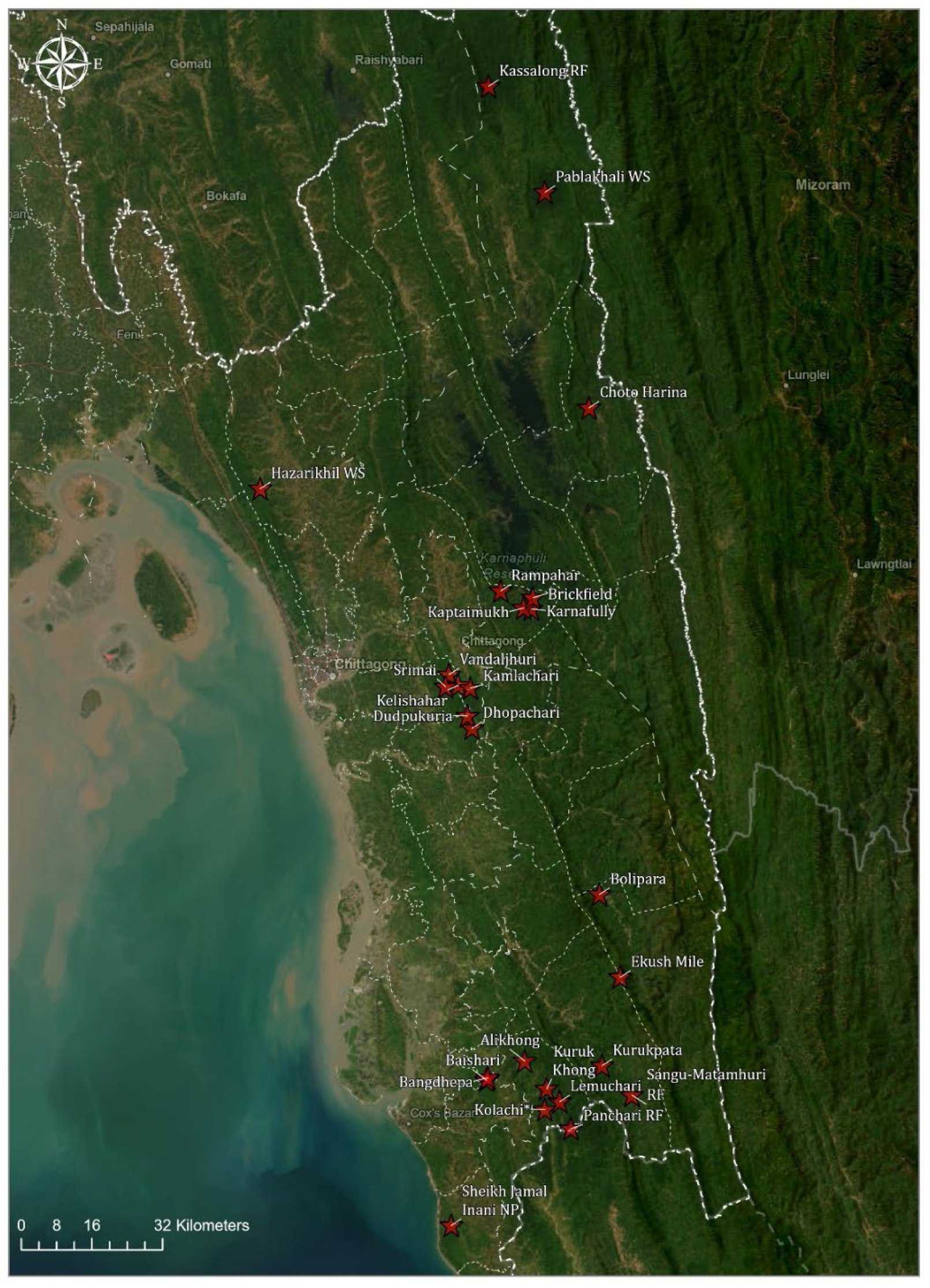
Locations of gibbon habitats in southeast Bangladesh

**Figure 2:**
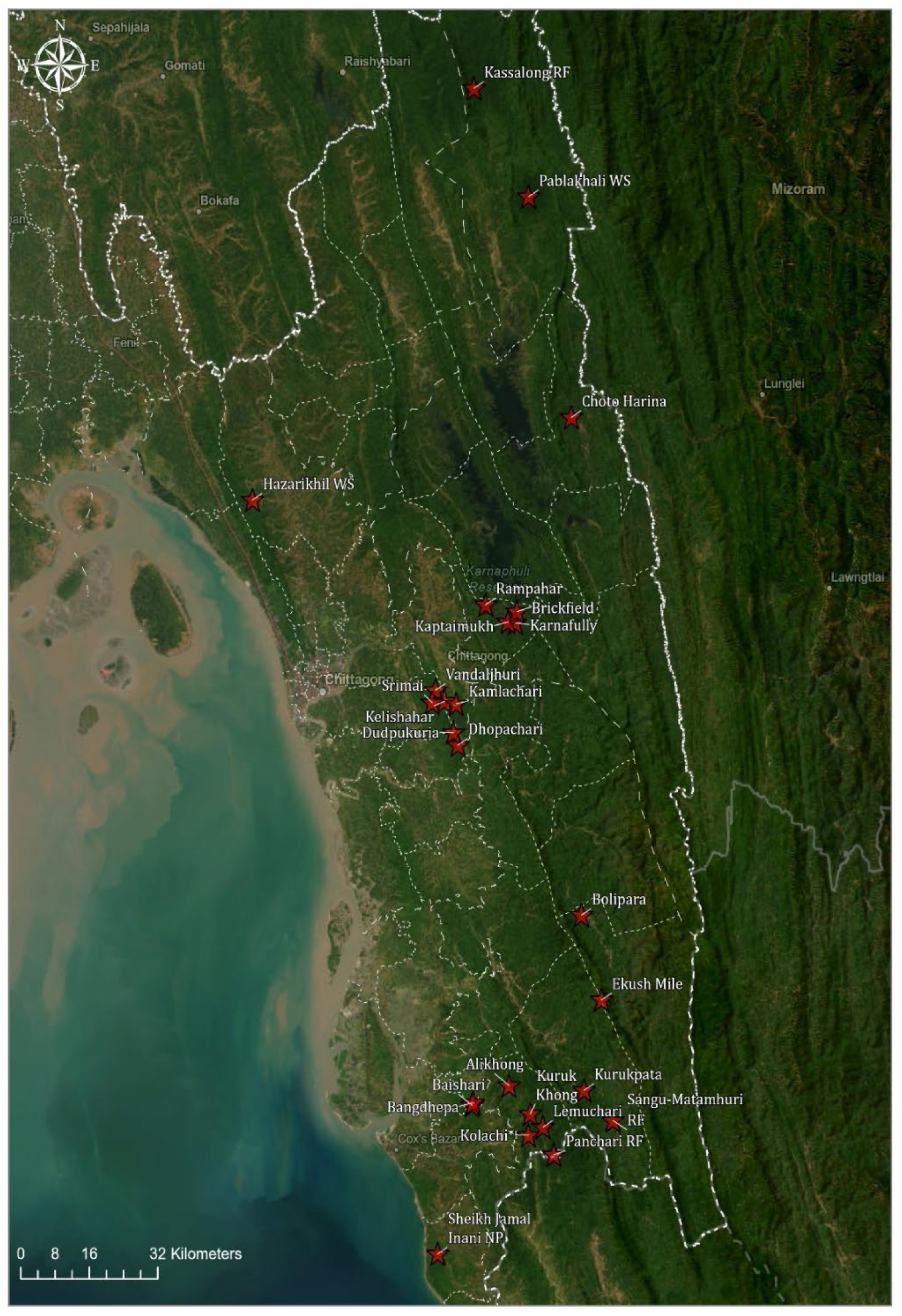
Locations of gibbon habitats in northeast Bangladesh

**Figure 3:**
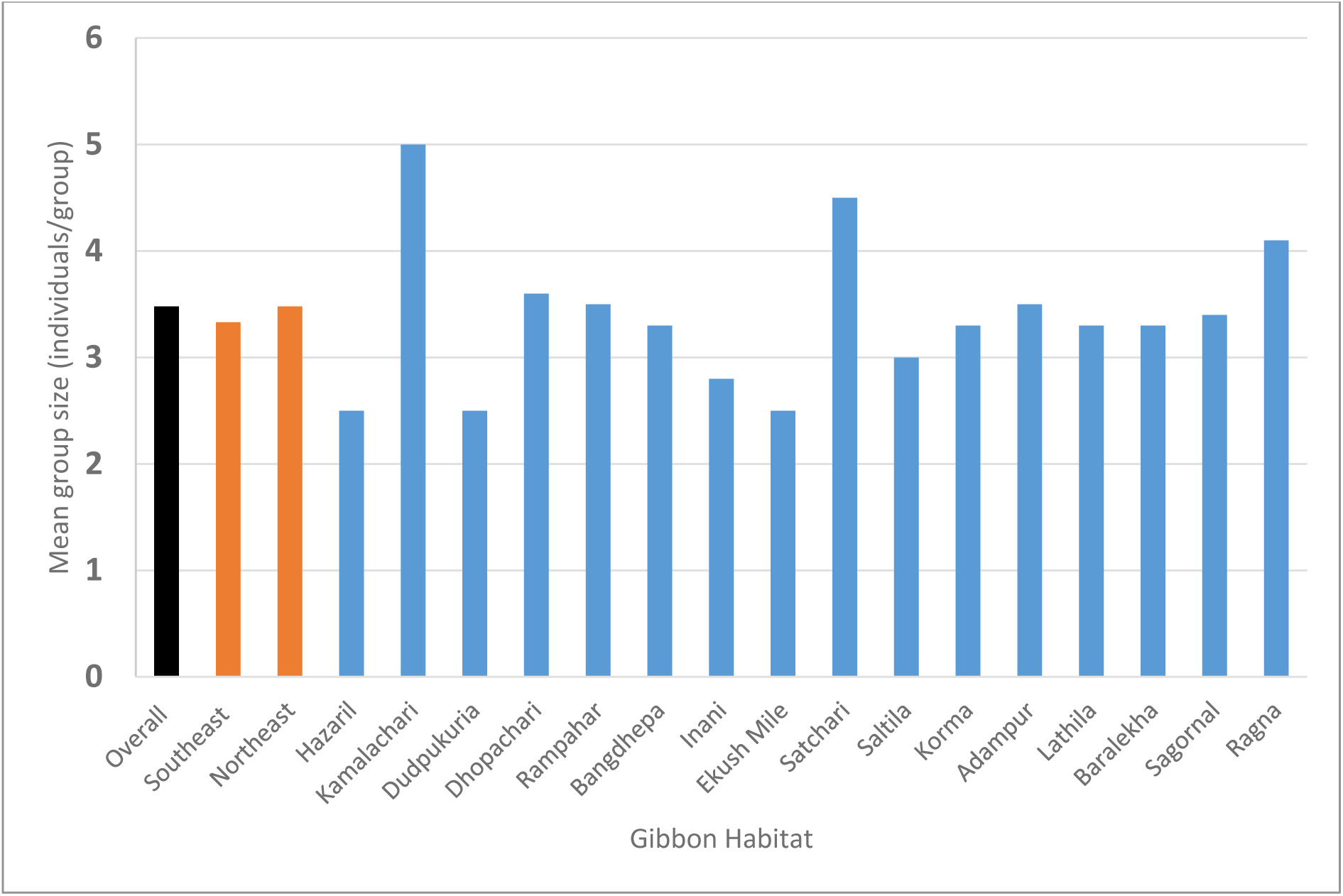
Groups sizes for western hoolock gibbons in Bangladesh (“overall”, indicated in black), northeast and southeast Bangladesh (regional averages indicated in orange), and for specific habitats.

Group sizes were also lower in habitats with areas <200 ha (n = 11, mean ± SD = 2.85 ± 0.898, range = 2 to 5).

Population density calculated as a proportion of total area for the 40 gibbon habitats for which densities could be estimated ranged from 0.01 to 7.69 groups/km^2^ (n = 40; mean ± SD = 1.20±1.49; 25^th^ percentile = 0.44; 75^th^ percentile = 1.2). Population densities were highest in three small and isolated areas: Kalapani (total area = 0.13 km^2^, density =7.69 groups/km^2^), Diari Biripata (total area = 0.6 km^2^, density = 3.33 groups/km^2^), and Srimai (total area = 0.29 km^2^, density = 3.45 groups/km^2^). If these three habitats are excluded, mean ± SD density was 0.91±0.931 groups/km^2^.

Population densities calculated considering only the portion of each gibbon habitat that was covered with forest ranged from 0.02 to 3.83 groups/km^2^ (n = 23; mean ± SD = 0.60±0.81), but most habitats had very low group densities, even if only forested area was considered (25^th^ percentile = 0.09; 75^th^ percentile = 0.77). Only Kaptai National Park (forested area = 15.47 km^2^, density = 1.03 groups/km^2^), Diari Biripata (forested area = 0.52 km^2^; density = 3.83 groups/km^2^) and Kolachi (forested area = 0.65 km^2^; density = 1.54 groups/km^2^) supported more than 1 groups/km^2^.

Our threat matrix analysis suggested that gibbon habitats in southeast Bangladesh experienced higher threats than habitats in northeast Bangladesh based on threat matrix scores. Our analysis rated 45.6% of habitats (n = 21) as highly threatened (threat scores = 15 to 20 on a scale of 3 to 20), 36.9% habitats (n = 17) were categorized as medium threatened (threat scores = 9 to 14) and 17.4% habitats (n = 8) are recognized as slightly threatened habitats (threat scores 5 to 8) (Tables 1 and 2). No habitat had a threat score lower than five, indicating that every remaining gibbon habitat in Bangladesh is facing anthropogenic threats.

## Discussion

This study used information from local community informants and other sources to identify 24 previously undocumented forest fragments occupied by western hoolock gibbons in Bangladesh, including 15 in southeast Bangladesh and 9 in northeast Bangladesh. Occupancy of each of these habitats was confirmed through extensive survey work. Our results greatly increase the number of habitats known to be occupied by gibbons, resulting in substantially higher estimates of the total population of gibbons in Bangladesh than those reported in recent literature (Islam et al. 2006, Naher et al. 2021).

Naher et al. (2021) identified Patharia Hill Reserved Forest, Rajkhandi Reserved Forest and Lawachara National Park in northeast Bangladesh and Kaptai National Park and Sangu- Matamuhuri Reserved Forest in southeast Bangladesh as the most important remaining gibbon habitats in Bangladesh, while Islam et al. (2006) recognized Adampur, Lathitila, and Lawachara National Park in northeast Bangladesh and Kaptai National Park in the southeast as important gibbon strongholds. This study confirms the importance of Lawachara National Park, Patharia Hill Reserved Forest (Baralekha, Lathitila), and Rajkandi Reserved Forest (Korma, Adampur, Kamarchara) in northeast Bangladesh, and adds Atora Hill Reserved Forest (Ragna and Sagornal), and Harinchara Tea Estate to the list of key gibbon habitats in that region. In the southeast, we confirmed the importance of Kaptai National Park and Sangu-Matamuhuri RF, and identify Sheikh Jamal Inani National Park, Baishari-Bangdhepa Wildlife Sanctuary, Dudpukuria- Dhopachari Wildlife Sanctuary, Kelishahar-Vandaljhuri forests, and Kassalong RF in southeast Bangladesh as additional important gibbon habitats. Naher et al. (2021) estimated that 76 gibbon individuals remain in Sangu-Matamuhuri RF. Sangu-Matamuhuri RF and Kassalong RF are the largest mixed-evergreen hill forest tracts in eastern Bangladesh and may support the largest population of the gibbons in Bangladesh. Further extensive population surveys, either through auditory sampling or passive acoustic methods may be helpful in better assessing the current status of gibbons of Kassalong RF and Sangu-Matamuhuri RF if the security situation does not allow for total population counts.

Naher et al. (2021) also reported the occurrence of gibbons at Fashiakhali Wildlife Sanctuary and Teknaf Wildlife Sanctuary in southeast Bangladesh but we did not find any signs of gibbons during our surveys there, which may indicate recent local extinctions at those sites. We suggest that further field investigations may be needed to confirm their status at Teknaf Wildlife Sanctuary. Conversely, Molur et al. (2005) suggested that gibbons had been extirpated from Harinchara Tea Estate and Hazarikhil Wildlife Sanctuary but we were able to confirm the persistence of gibbons in those sites.

Observed population densities of western hoolock gibbons globally range from 0.3 to 4.4 group/km^2^ (Green, 1977; Gittins, 1980). Chowdhury et al. (2009) estimated the population density in Assam, India, as 2.43 groups/km^2^ and Ray et al. (2015) estimated mean density of 3.65 groups/km^2^ from Namdhapa National Park of Northeast India. Tilson (1979) estimated a group density of 0.55groups/km^2^ in the Indian state of Tripura (Mukherjee, 1982) and Alfred and Sati (1990) reported a density of 3.01groups/km^2^ from West Garo Hills. Previous studies in Bangladesh have reported gibbon densities on the low end of the global range, including an estimate of 0.3 group/km^2^ or 0.23 group/km^2^ in Rajkhandi forest (Siddiqi, 1986) and 1.12 groups/km^2^ at Lawachara National Park (Osterberg, 2006). More recently, Naher et al. (2021) estimated gibbon population densities between 1 and 2 groups/km^2^ at Satchari (1.12 groups/km^2^),

Lawachara (1.69 groups/km^2^), Rajkhandi (1.27groups/km^2^), Patharia (1.19 groups/km^2^), Kaptai NP (1.11 groups/km^2^), and Sangu-Matamuhuri RF (0.72 groups/km^2^). Most of our population density estimates were quite low compared with western hoolock gibbon densities reported in other range countries, and lower than densities reported in earlier studies in Bangladesh.

Bangladesh is among the most densely populated nations in the world, and has the highest population density among countries with areas >10,000 km^2^ (https://www.worldometers.info/world-population/bangladesh-population/). Anthropogenic pressure on gibbon habitats has led to habitat loss and extensive anthropogenic disturbance inside forested areas in Bangladesh. As a result, western hoolock gibbon populations in Bangladesh are highly fragmented and this globally endangered species is believed to be more critically threatened in Bangladesh than in the other range countries of India and Myanmar (Molur et al., 2005).

To the best of our knowledge, our study is the first to intensively engage local communities in locating a wildlife species in Bangladesh. Local communities near gibbon habitats regularly encounter gibbons or hear their calls. Due to their loud and unique songs and their distinctive locomotor behavior involving brachiation, gibbons are unique among primates and can easily be recognized by non-scientists (Ahsan, 1994). Therefore, local communities often hold knowledge that can be engaged to identify gibbon habitats, especially in areas that are remote and difficult to access for non-residents. Citizen science has been used to assess the populations of many animals worldwide including Raffles’ banded langurs (*Presbytis femoralis*) in Singapore (Ang et al., 2021), Pallas’s fish eagle (*Haliaeetus leucoryphus*) in Bangladesh (Chowdhury et al., 2022), slender lorises in India (Bhaskaran et al., 2015), hornbills (Yeap and Perumal, 2017) in Peninsular Malaysia, orangutans in Malaysian Borneo (Oram et al., 2022 and three species of small apes in Peninsular Malaysia Mohd Rameli et al. (2019). Our results confirm that community consultation can be an important tool for identifying western hoolock gibbon populations, estimating their population sizes, and identifying specific threats affecting local landscapes in Bangladesh.

The identification of several new western hoolock gibbon habitats and higher estimate of the total gibbon population in Bangladesh should not be misunderstood as suggesting an actual increase in the gibbon population in Bangladesh. We identified additional populations of gibbons through comprehensive and successful engagement with the local communities and through extensive long-term fieldwork over a larger area than previous studies, which allowed us to identify populations that had not been previously surveyed. Nonetheless, habitat quality in the eastern hill forests in Bangladesh is rapidly declining (FD, 2020), and the actual population that can be supported by these habitats is almost certainly decreasing.

The mean group size of studied western hoolock gibbons in Bangladesh appears to be lower in smaller habitat fragments, which suggests that these populations may not be reproducing at rates adequate to maintain their population sizes in the long term, a pattern that has also been reported in India. For example, Kakati et al. (2009) recorded the mean group size of 2.5 (n = 2) for populations in small forest fragments, 3.29 (n = 24) in medium-sized fragments, and 3.9 (n = 28 groups) in larger forest fragments. Some previous studies also suggest local population declines at specific sites from which multiple assessments are available. At Lawachara National Park, mean group size was recorded as 3.5 (n = 6) in 1984 (Gittins and Tilson, 1984), 3.17 (n = 6) in 1990-1991 study period (Feeroz and Islam, 1992), and 3.0 (n=8) in 1989- 990( Ahsan, 2001).

However, our study estimated a mean group size of 3.42 (n = 8) at Lawachara National Park, suggesting that the population size at Lawachara may have remained relatively stable since 1984, despite some fluctuation.

The western hoolock gibbon is legally protected in Bangladesh by the Wildlife (Conservation and Security) Act 2012, Schedule I. Hunting, trapping, and shooting of gibbons are strictly prohibited by law. However, all gibbon populations and their habitats face severe threats exacerbating the risk of rapid local extinctions. Therefore, the following conservation initiatives should be initiated to protect the remaining western hoolock gibbons of Bangladesh:

- Laws against hunting, encroachment, and habitat destruction should be strictly enforced, and local communities should be engaged to raise conservation awareness about legal prohibitions and the negative impacts of gibbon hunting and illegal trade;
- Involvement of the Khasia communities for gibbon conservation to prevent further extensions of pan punji (betel leaf vineyard plantation) and reduce existing threats to the gibbons;
- Further extension of orchard plantations (i.e., lemon, banana etc.) should be monitored and strictly discouraged inside gibbon habitats;
- Firewood collection for brickfields should be monitored and strictly discouraged within gibbon habitats;
- Isolated gibbon habitats should be connected by fast-growing native tree species and habitat restoration programs should be initiated in degraded habitats using native fruiting trees that provide gibbon foods;
- Monoculture plantations should be discouraged inside the gibbons habitats;
- Co-management Committees should be engaged to co-formulate a long-term conservation plan to reduce threats to the forests and gibbons habitats;
- Community-based long-term and sustainable conservation strategic plans should be developed and implemented; and
- Communication, education, and public awareness programs should be implemented to develop positive behaviors and attitudes among relevant stakeholders to support sustainable gibbon conservation.

## ACKNOWLEDGMENT

We grateful for the financial support from the Bangladesh Forest Department through their SUFAL Innovation Grant, IUCN Section on Small Apes, Wildlife Conservation Network, Gibbons Conservation Alliance, International Primatological Society, American Society of Primatologists, Primate Conservation Inc. Primate Conservation Trust, Primate Action Fund and Rufford Foundation Small Grant through the Malaysian Primatological Society. We thank the Bangladesh Forest Department for permitting us to carry out the fieldwork in the eastern hill forests of Bangladesh. We also acknowledge to Dr. Susan Cheyne, Dr. M Farid Ahsan, and Mr. Ishtiaq Uddin Ahmad for their continuous encouragement and support to carry out this work. We are grateful to School of Biological Sciences, Universiti Sains Malaysia for providing the necessary support. This work would not have been possible without the generous support from the numerous local field assistants - we owe you.

